# Label-free virus-antibody interaction monitoring in real time by common-path interferometry

**DOI:** 10.1101/2022.08.23.504719

**Authors:** Samer Alhaddad, Houda Bey, Olivier Thouvenin, Pascale Boulanger, Claude Boccara, Ignacio Izeddin, Martine Boccara

## Abstract

Viruses can affect all life forms, raising concerns about virus detection and quantification of small nanoparticles. In this paper we use a label-free, full-field, incoherently illuminated common path interferometric method to detect, track, and quantify biotic nanoparticles. The detection consists of amplifying the light scattered by single nanoparticles in the sample solution. Then, the use of single-particle tracking analysis is used to monitor the change in particle diffusive mobility. With this approach, the recognition signature of T5 phages with purified antibodies targeting the major capsid protein is detected in a few minutes. We also tracked the interaction between SPP1 phages and physiological non-purified serum-containing multiples antibodies molecules. The first interactions occur after around one minute, and the recognition signature is detectable after minutes. In addition, we have been able to differentiate two populations of similar size of empty and full (encapsulating DNA) capsids of T5 in a heterogeneous solution demonstrating the robustness of this label-free detection approach. Furthermore, by combining the diffusion coefficient to the number of tracked particles, we were able to estimate the affinity of the virus-antibodies reaction.

## 1. INTRODUCTION

The ability to detect and characterize viruses rapidly and efficiently plays a crucial role in avoiding global widespread of severe health issues. In particular, early detection of viruses helps provide targeted therapy, extends the treatment space, reduces morbidity, and limits the transmission of viruses. Therefore, there has been considerable interest in nanoparticle detection, especially in the biomedical and pharmaceutical fields as well as in virus research by improving testing and diagnostics.

Numerous existing methods for virus detection can broadly be divided into three categories. Firstly, techniques based on measuring the infectivity include virus culture in eukaryotic cell lines or bacteria. These cells represent a suitable host to be infected by the virus. An appropriate incubation period results in the formation of plaques, which can be fixed and stained for visualization. Plaque assay is the standard gold method for quantifying infectious virus titer and can take up to several days to obtain the results [1]. Secondly, techniques that consist of examining the viral nucleic acid or protein of viruses include PCR (Polymerase Chain Reaction) amplification techniques or antigen-based tests. PCR techniques are known for their specificity, and they consist on searching for the presence of specific genetic material in the virus, and then amplifying a genetic sequence of interest to quantify its presence in the initial sample, taking up to a few hours to get the result [2]. Antigen-based tests are used to detect specific viruses, and these tests look for proteins on the surface of the virus. Results can be obtained quickly in tens of minutes [4] despite a major issue of lack of sensitivity [5]. The third category relies on a direct count of virus particles such as viral flow cytometry which is based on counting individual particles in the solution sample. Dynamic light scattering method is also used to estimate the ensemble average size of particles in solutions based on the diffusion coefficient of particles in the sample solution. Electronic microscopy techniques have been widely used to study the detailed morphology of viruses in terms of structure and composition [3].

Light microscopy technique is a powerful tool to study biological samples, for its non-invasiveness and potential high throughput. Nanoparticles and virus particles are too small to be visualized directly with conventional optical microscopy due to their weakly scattering characteristics resulting from their small size and low refractive index difference compared to its surrounding medium. Overall, the scattered intensity induced by the illumination of the particles scales with the sixth power of the particle size, which makes the particle indistinguishable from the background illumination due to the lack of contrast. In order to overcome the lack of contrast, fluorescence microscopy methods are usually employed, where the targeted molecules are labeled with fluorescent labels and therefore detect the molecule of interest without getting affected by the non-labeled background. For virus particles, fluorescent labels have been mostly used to tag the capsid protein [6] or to tag DNA viral genomes [7]. To note that a rapid virus detection method has been reported requiring about one minute to obtain a result, in which calcium labeling has been used to track the mobility of viruses when interacting with specific antibodies [8]. However, the optimization of the labeling process is crucial in order to retrieve high contrast images and yet, fluorescence labels suffer from phototoxicity, photobleaching, photostability, and saturation.

Highly sensitive optical interferometric microscopy techniques have changed the ability of optical microscopy due to its conception based on signal amplification using the interferometry principle. Although the intensity scattered from small particles is very weak compared to the illumination light, the concept of interference makes the detection of nano-sized particles achievable with a camera-based detector. Different interferometric microscopy configurations have been used to detect, identify and track small particles, molecules, metallic particles, and other materials in a wide range of domains [9, 10, 34]. These imaging methods have been applied to study dynamics in living cells [11, 12], characterize single particles and molecules in solutions [13, 14], as well as sensing and imaging proteins interactions sites applications [15, 16]. However, the main drawback of label-free interferometric methods is the lack of specificity. Many groups tackled the specificity challenge in different ways. Functionalizing surfaces and nonmaterial provide a means for targeting through selective binding to receptors at targeted sites [11-15]. Other groups studied the interaction between proteins by monitoring the interferometric contrast modification resulting from an agglomeration of molecules [23]. These interferometric scattering methods proved their sensitivity by detecting single protein and their sensibility by the ability of distinction between different agglomerations [31]. More recently, commercial interferometric scattering-based instruments have been used to characterize the interaction between antibodies and spike protein of virus of interest [32].

We have previously addressed the question of detecting, tracking, and counting viruses and vesicles in an aquatic environment. As a solution, we developed a full field incoherent illuminated common path interferometer working in transmission [17, 18]. With this approach, the magnitude of the interferometric signal has been used in complement to single particle tracking analysis in order to distinguish between different populations of particles of similar size and different nature. In this work, we have been able to improve the signal-to-noise ratio of our set-up using a high-speed camera with a high full well capacity, allowing a better estimation of the dynamics of the particles. Moreover, we introduce a new approach to virus detection by adding specific antibodies to the simple solution containing the targeted virus. By these means, we monitor the interaction between viruses and antibodies molecules in solution in terms of variation of interferometric signal contrast and diffusion mobility of particles. This approach is demonstrated using purified IgG molecules anti pb8 with T5 phages and a non-purified serum-containing multiple antibodies anti-SPP1 and SPP1 phages. Despite the heterogeneity of the particles in a non-purified serum, we could detect a recognition signature of the antibodies reaction with antigens present on the virus’s outer surface in about 1 minute, and the aggregation is detected in a few minutes. By relying on the variation of the number of the tracked particles in the solution, we were able to estimate the molecular dissociation rate of the antibodies in use.

## 2. METHODS and MATERIALS

Interferometric microscopy has proved its potential as a powerful bio-imaging method. In this study, we use a similar set-up for the common path interferometer as described in [17]. In interferometric scattering microscopy working in transmission, the camera performance is critical. For this reason, in this work, we chose a new sensor and an improved full well capacity (FWC) combined with a much faster acquisition leading to a significant improvement of the signal-to-noise ratio (SNR). This allows the detection and tracking of small moving scatterers with a better precision.

### 2.1 Principles of the virus detection and interferometric signal

The modality used through this work consists of measuring the interference signal between the reference field coming from the light source and the scattered field of the nanoparticles moving in the liquid sample. The intensity recorded on a camera pixel results from the sum of the reference beam and light immerging from the nanoparticles. It can be written as:

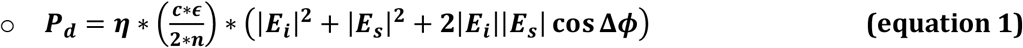

Where ***P***_***d***_ represents the power intensity recorded on the camera, ***η*** is the camera quantum efficiency, ***c*** is the celerity of light in the free space, ***ϵ*** is the permittivity of the medium, ***n*** is the refractive index of the medium, ***E***_***i***_ is the incident field, ***E***_***s***_ is the scattered electric field and **Δ*ϕ*** represents the phase difference between the reference electric field and the electric field extinction of the nanoparticles.

In transmission, the phase difference designates mainly the Gouy and defocus phase shift that is caused by the variations of the wave vectors at the focal plane of the objective lens and the phase contribution determined by the dielectric function of the nano-object [9]. For small particles such as viruses; the phase can be neglected. Therefore, constructive and destructive interferences occur depending on the axial position of the particle moving in the solution. The scattered field emerging from the particle is proportional to the incident reference field:

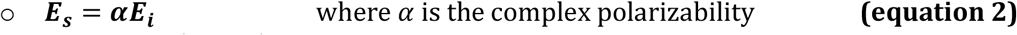

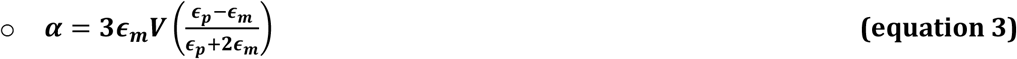

### (Equation 3) is applicable in the Rayleigh approximation regime (diameter << wavelength)

Here ***ϵ***_***m***_and ***ϵ***_***p***_are the permittivities of the surrounding medium and the particle respectively. Those wavelength dependent complex quantities determine the balance between absorption and scattering contribution at a specific wavelength. ***V*** designates the volume of the particle.

Returning to the recorded power intensity ***P***_***d***_, the static background coming from the LED illumination is dominant over the interference and scattered terms. For small particles, the scattered field is negligible compared to the interference term representing the dynamics of the particles in the solution.

A schematic drawing of the full field common path interferometer is shown in **Fig1** (a). Illumination of the sample solution is provided by a 455 nm light-emitting diode (LED). Light pass through the solution containing nanoparticles, a water immersion objective lens 100X/1 NA (Olympus, Japan) is used for collection and imaging purpose; the beam is then directed to tube lens (300mm, Thorlabs, USA) to focus the light on the camera. The interferometric signal and the diffusion properties depend on the size of the particles and the refractive index difference between the particle and its surrounding medium. Thus, these two parameters are used to distinguish between particles of similar size but with different internal components in a first approach and to follow reaction between viruses and specific antibodies, **Fig1**(b).

**Fig 1:**
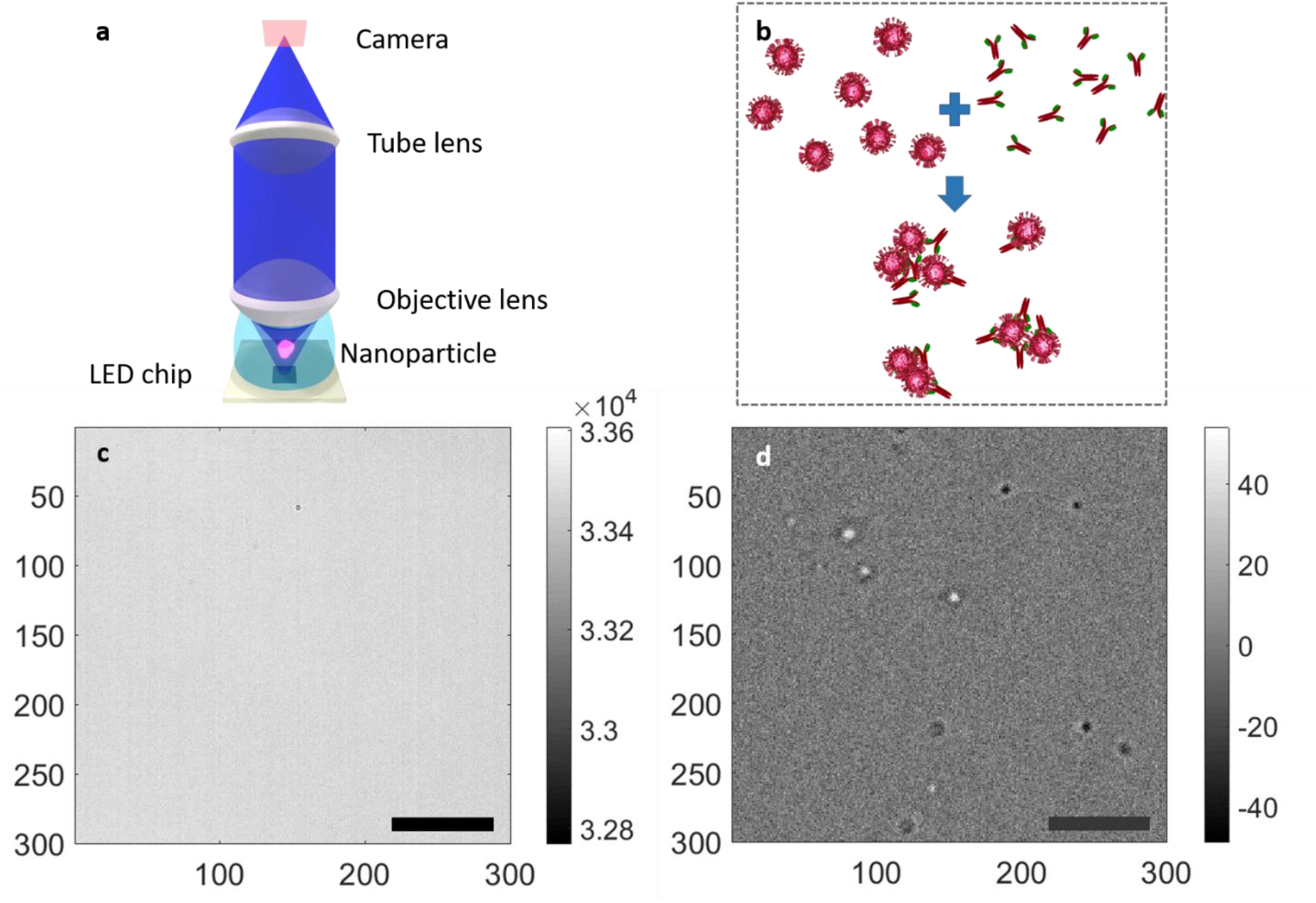
(a) Common path interferometer description: an incoherent 455nm LED illuminates the sample with a full angle of 80 degrees (Thorlabs M455L3). Light travels through the sample solution, light coming from the LED and light scattered from the nanoparticles is collected using a water immersion objective lens (Olympus 100x, NA=1) and focused using a tube lens (focal lens 300mm) on an ADIMEC Quartz 2A750 camera. (b) A crude illustration of specific antibodies-viruses reaction followed by the aggregation of particles. (c) Zoom on a direct image before processing frame, scale bar: 5µm. Images are acquired at 500 FPS with an exposure time of 1.8ms. (d) Processed image, in which the stack of images are normalized to 4000 and the average frame of the entire stack is subtracted from every frame image, scale bar: 5µm. Alternating bright and dark spots represent the image of 100 nm in diameter particles immersed in water.

### 2.2 improving signal to noise ratio and acquisition speed

A major new modification in our setup is the introduction of a new camera (Quartz 2A750, ADIMEC, Netherlands) that has a maximum frame rate of 720 frames/s at 1440×1440 pixels and a full well capacity of **2** * **10**^**6**^electrons. This new camera results in an improvement of the SNR, about three times for the shot noise level compared to our optical set-up described in [17]. Here, images are acquired at a speed of 500 frame/s with a field of view of 103*x*103 *µm*^2^. A typical acquisition consists of 500 images.

### 2.3 Single particle tracking and interferometric signal

We first remove the static background, from raw images in **fig 1**(c), by subtracting the temporal average frame value of the 500 frames stack from every image frame. The resulting images are similar to those obtained by single molecule fluorescence microscopy images, with the difference that now, the signal intensity of the point of spread function (PSF) contains useful information. The bright and dark spots on the image indicate thus constructive and destructive interferences (fig 1(d)).

For single particle tracking, we have used a modified version of the multiple-target tracing (MTT) algorithm, which was initially developed for fluorescence applications [19].

In **fig** 2, we show the analysis of particle tracking using nanoparticles of *Sio*_2_of 100nm in diameter, with a concentration of 10^9^ *particles/mL*. Fig2 (a) represents the mean squared displacement plot of individual particles. Here we count 806 detected particles for a single acquisition of 500 frames. We then associate the maximum absolute value of the interferometric signal per particle to each trajectory because the interferometric signal of each particle varies with the changes of the axial position of the particle. We ensure that particles moving inside the solution will pass by a maximum of intensity characterizing its scattering behavior in the volume of detection. Fig2 (b) represents the histogram of the interferometric signal of all the particles; a homogenous normal distribution can be seen with an average value of 108.7(a.u.) with a standard deviation of 9.61(a.u.). The interferometric signal depends on the size and the refractive index of the particle. Finally, we perform a single particle analysis in order to evaluate the diffusion coefficient of the nanoparticles.

**Fig 2:**
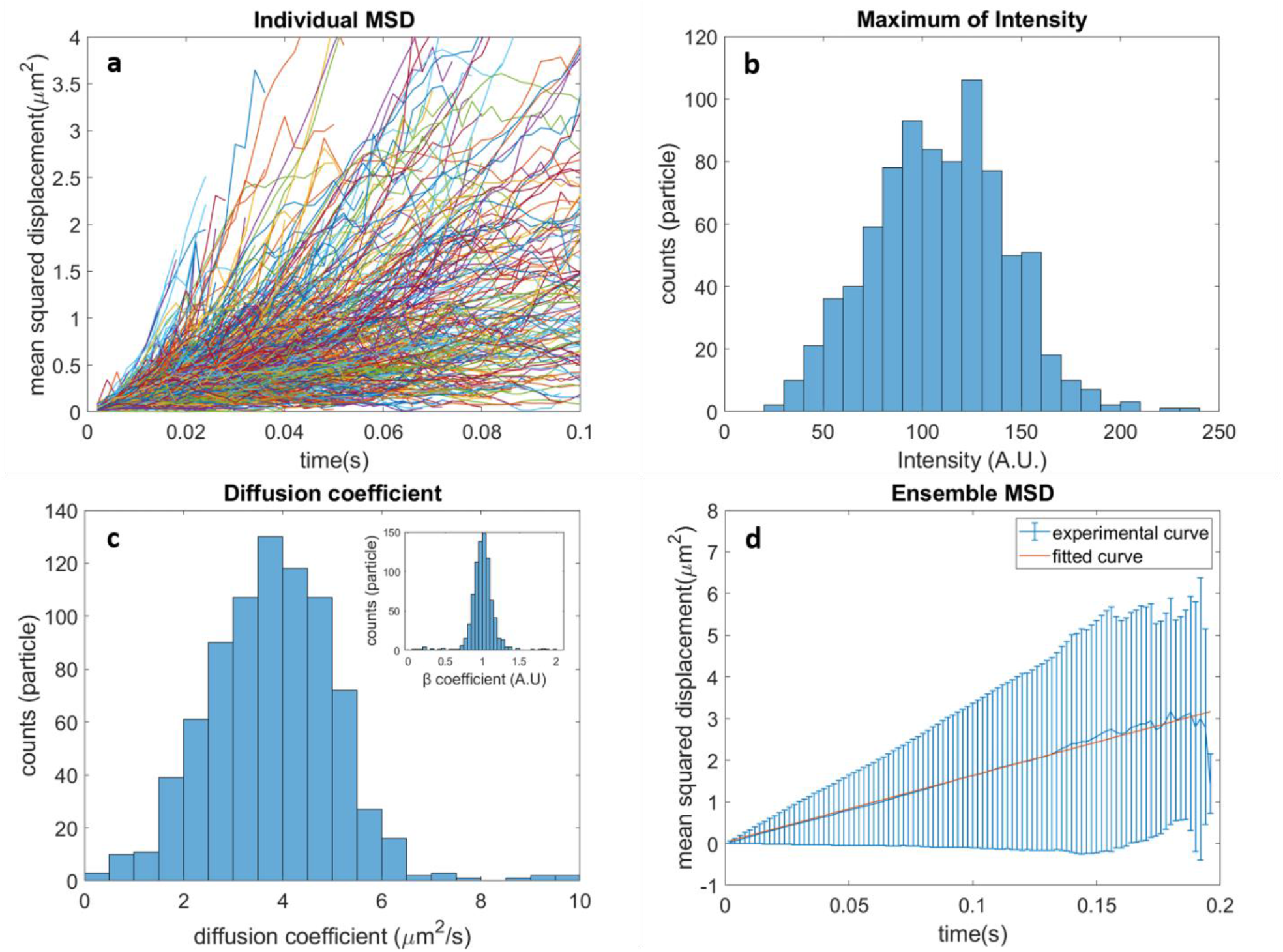
Validation of the single particle analysis using 100 nm diameter *SiO*_*2*_ particles (a) Individual mean squared displacement (MSD) curves of particle trajectories. (b) Histogram of distribution of the absolute maximum of intensity per trajectory, average estimated value 108.7±9.61 A.U. (c) Histogram of the fitted diffusion coefficient resulting from individual MSD; average estimated value 3.82 ±1.01*µm*^2^*/s*. The inset histogram represents the fitted beta coefficient with an average of 1.009 ±0.019. (d) Plot of the ensemble average MSD of all the trajectories and the linear fitting of the MSD, average fitted diffusion coefficient the second and the first 20% of the data points is 3.85*µm*^2^/*s*. The data results from 806 tracked trajectories, 500 frames acquisitions at 500 frames/s.

To estimate the individual diffusion coefficient and characterize the diffusion regime of each particle, we fit the MSD plot presented in **Fig2** (a). The mean squared displacement can be expressed as:

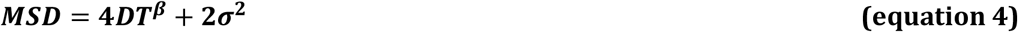

Where ***D*** represents the diffusion coefficient, ***T*** is the lag of time between images, ***β*** indicates the diffusion regime of the particle and ***2σ***^**2**^ is the mean localization error in x and y.

***β*** is one for normal Brownian diffusion; this term is used to examine the deviation of the particle from normal diffusion, ***β*** >1 indicates a super diffusion behavior and ***β*** <1 is a sub diffusion signature. We apply an unweighted linear regression between the second point and the first 20% of the MSD curve since the nanoparticles are immersed in an isotropic water-like solution [20-21]. If we assume that the particles move randomly (***β*** = **1**), the estimated diffusion coefficient of *Sio*_2_of 100nm in diameter in **Fig2** (c) shows the histogram of individual diffusion coefficient values centered at 3.82 ±1.01*µm*^2^/*s*. Similarly, the inset plot in **Fig2**(c) confirms the Brownian diffusion of the particles, ***β***=1.009 ±0.019.

In addition to the trajectories analysis, an ensemble average analysis can be employed if we assume a homogeneous distribution of the particle size. In **Fig2** (d), we have estimated a diffusion coefficient of 3.85 *µm*^2^/*s* for particles of *Sio*_2_of 100nm in diameter when fitting linearly the ensemble average MSD. This value is similar to the one found with individual trajectory analysis, indicating the homogeneity of the solution.

### 2.4 Estimation of the molecular affinity of antibodies using single particles tracking analysis

The binding forces between specific antibodies and targeted protein depend on molecular properties of the interacting species. On a molecular scale, the dissociation constant can be described by the ratio using the concentration of products and reagents, at the equilibrium state, (equ 5,a,b). The rate equation can be estimated in a straightforward way (equ 5,c).

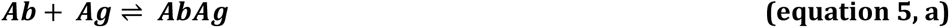

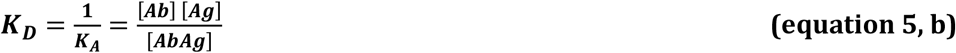

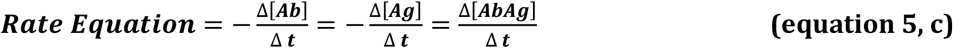

Where ***Ab, Ag, AbAg*** represent antibodies molecules, antigen molecules and the recombination of both molecules. ***K***_***A***_, ***K***_***D***_ are the association and dissociation constants respectively

On the macromolecular scale, the reaction gets more complicated because of the various interaction between viruses in the solution in presence of antibodies. The description of the reaction defined by consecutive avalanche reaction as involves the concentration of all species for each step of the reaction, as can be seen in (equ6).

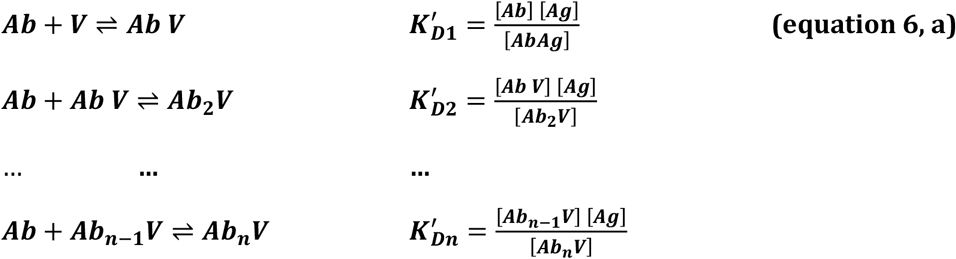

***V*** represents the virus particle, 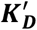 is the dissociation constant for each step. ***n*** is the number of binding sites step present on the external surface of the virus.

Therefore, a saturation function can be used to describe when the system reaches an equilibrium state (equ 6,b). **r** defined as the ratio of bound particles over the initial concentration of virus in the solution sample [33]. Therefore, the **Kd** value of the interaction can be estimated for specific antibodies using the concentration value of bound antibodies and the number of possible binding sites **n** per virus capsid.

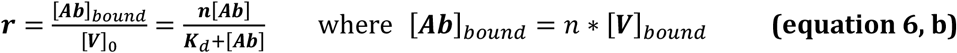

On a higher scale, the interaction between viruses and antibodies can be represented as:

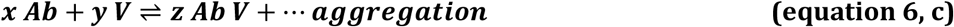

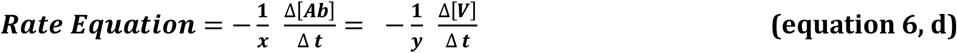

Where ***x, y, z*** are the stoichiometric coefficients of the reaction for each species: antibodies, viruses and aggregates virus and antibodies respectively.

The concentration of bound antibodies can be estimated using the number of particles that did not interact with other particles until the system reaches its equilibrium (**equ6**,**b**). In practice, we can retrieve the number of these particles using the variation of their diffusion coefficient when adding antibodies, ***V***_*bound*_ = ***V***_0_ − V(*t*). Here, we employed the number of non-interacting particles since the aggregation process is a heterogeneous process.

### 2.5 Phages, capsids and antibodies

Purified T5 phage and SPP1 phage were used in this study. The initial concentration of T5 bacteriophages was estimated to 7.3 ×10^12^

PFU/mL (PFU: plaque forming unit). Full (with DNA) and empty capsids of T5 were also analyzed with our set-up. Full capsids, have an initial concentration of 10^12^ heads/mL [24] and empty capsids, have an initial concentration of 0.6 mg/mL (about 4*10^13^ heads/mL). Phages and capsids were diluted in T5 buffer (10mM Tris-Cl pH=7.4, 100mM NaCl, 1mM *MgCl*_2_). SPP1 bacteriophage has a morphology similar to T5, but capsid is ∼60 nm [25], has an initial concentration: is 2.24*10^11^PFU/mL. SPP1 was diluted in buffer (100mM Tris-Cl pH=7.5, 100mM NaCl, 10mM *MgCl*_2_)).

Purified polyclonal IgG against T5 major capsid protein (pb8; 800 sites/capsid) have an initial concentration of 0.84 mg of protein/mL. Serum anti all SPP1 particles [26-27], have also been used, we estimated 5 times more serum albumin compared to IgG molecules. We measured a concentration of 0.1mg/mL in the supernatant of anti SPP1 serum after centrifugation at 14000 rpm for 10 minutes to eliminate big aggregates.

## 3. RESULTS and DISCUSSION

### 3.1 Differentiation of empty versus full phage T5 capsids

In order to demonstrate that our detection approach using single particle (SPT) analysis is well suited to detect and differentiate NPs of similar size and of different composition, we choose to use capsids of T5 phage containing DNA (full capsids) and capsids of similar size lacking DNA (empty capsids). As expected, these populations of particles must have a close value of diffusion coefficient in a liquid solution. However, the refractive index of the full capsids is expected to be higher compared to empty capsid. Acquisitions were processed in order to retrieve the diffusion characteristics of the particles as well as their interferometric signal. We first estimate the diffusion coefficient and the interferometric intensity for each homogeneous solution of the two populations.

In terms of particle mobility, we estimated an individual diffusion coefficient of 3.92±0.87 *µm*^2^/*s* for 192 tracked empty T5-capsids compared to 4.05±0.98 *µm*^2^/*s* for 471 tracked full T5 capsids. **Fig 3**(a) represents in a boxplot presentation the distribution of the estimated diffusion coefficient for each sample. In addition, the diffusion coefficient resulting from the ensemble average MSD are represented in Fig3 (a), an ensemble diffusion coefficient of 3.86*µm*^2^/*s* and of 4.03*µm*^2^/*s* have been found for Empty and Full capsids respectively. A nominal concentration of 10^9^*particles/mL* was used for each acquisition.

**Fig 3:**
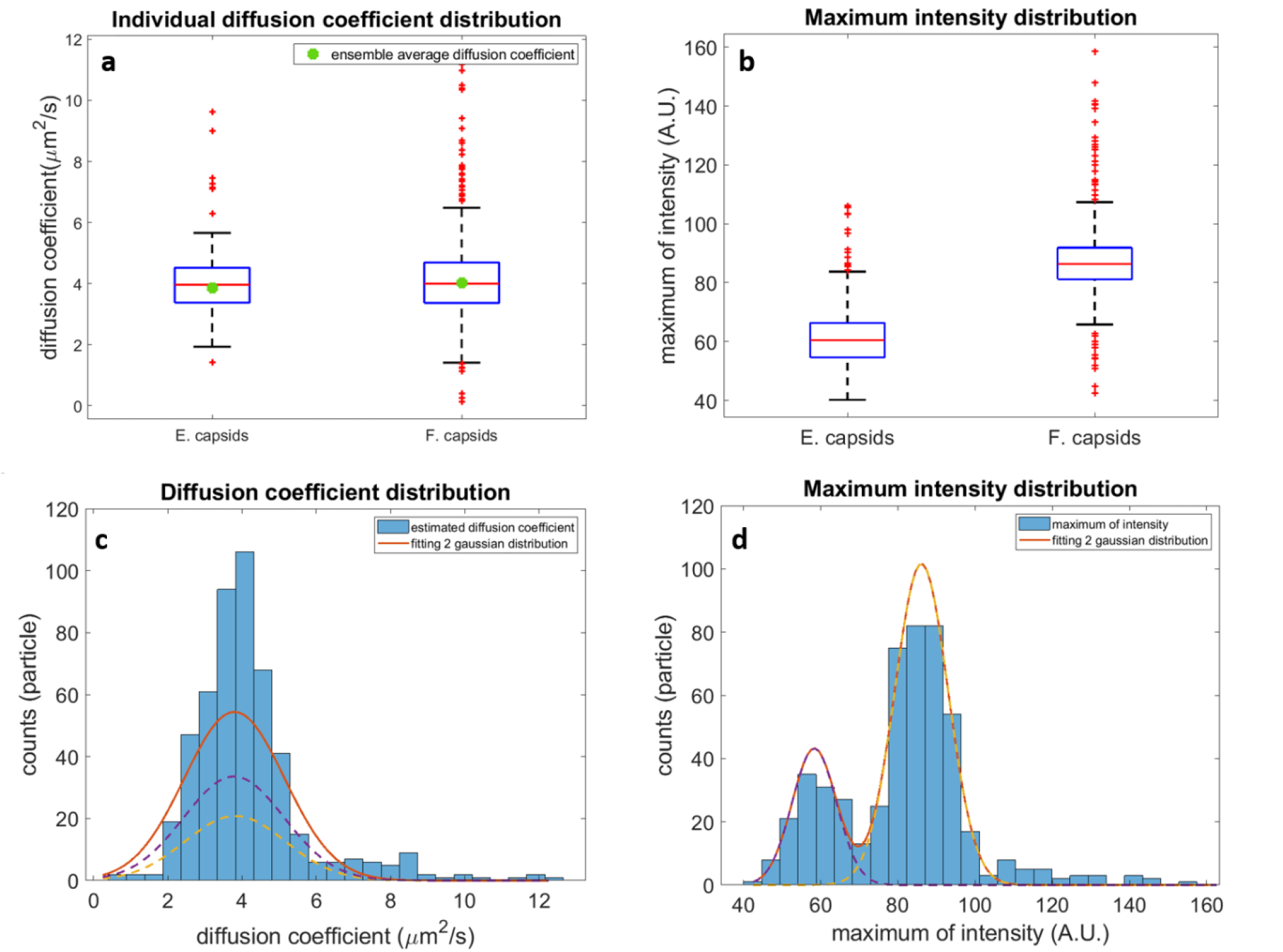
(a) boxplot represents the distribution of the individual diffusion coefficient for empty and full capsids. For empty capsids: the median =3.92 *µm*^2^/*s* and the 25th and 75th value are between 3.35 and 4.61 (lower and upper limit of the blue square), for full capsids: the median =4.05 *µm*^2^/*s* and the 25th and 75th values are between 3.48 and 4.75. The ensemble average diffusion coefficient when fitting the Ensemble average MSD, 3.86 *µm*^2^/*s* for Empty capsids and 4.03 *µm*^2^/*s* for Full capsids (b) boxplot represents the maximum of intensity distribution for empty and full capsids, for empty capsids: the median =60.49 *µm*^2^/*s* and the 25th and 75th value are between 54.57 and 66.28, for full capsids: the median =86.38 *µm*^2^/*s* and the 25th and 75th value are between 81.175 and 91.9. (c) Histogram of distribution of the estimated Diffusion of a mixture of Empty and Full capsids. The estimated two Gaussian normal distribution fitting has amplitudes of (20.85,33.57), mean values of (3.82, 3.77) and a standard deviation of (1.9, 1.9) respectively. (d) shows the histogram of maximum of intensity, The estimated two normal distribution fitting represents amplitudes of (101.63, 43.26), mean values of (88.03, 58.33) and a standard deviation of (9.75, 8.11) respectively.

Despite the similarity in terms of particle mobility, a difference of interferometric intensity has been detected for these samples. An average intensity of 60.49 ± 6.25(*A. U*.) is found for empty T5-capsids compared to a value of 86.38 ± 9.68 (*A. U*.) for full capsids, as shown in **Fig3** (b). These results clearly show that we are able to differentiate between full and empty capsids solutions using the interferometric signal despite the similarity in the diffusion properties. We noted a higher standard deviation when estimating the diffusion coefficient as well as the interferometric signal which indicates a larger dispersion of full capsids compared to empty capsids, this observation confirms the morphological study of the capsids using transmission electron microscopy [24].

In a second experiment, we used a heterogeneous solution containing both types of capsids to distinguish between the two populations of capsids by comparing their interferometric signal contrast. For this purpose, we used single particle analysis approach. We chose a similar concentration of empty and full capsids. As expected, we are not able to distinguish between the two populations in terms of diffusion coefficient as shown in **Fig3** (c). The fitting of the histogram of diffusion coefficient with two Gaussian distributions using least squared error fitting results in an overlap of the curves with a diffusion coefficient of 3.82 ± 1.9*µm*^2^/*s* and 3.77 ± 1.9*µm*^2^/*s*. However, when fitting the histogram of the interferometric signal with two Gaussian distributions, two distinct populations were found (**Fig3** (d)). A first population found at 58.33 (*A. U*.) with an amplitude of 43.26 and a standard deviation of 8.11 and a second population centered at 88.03 (*A. U*.) with a standard deviation of 9.75 has an amplitude of 101.63. The first population designates the population of empty capsids of T5 while the second population consists of full capsids that have higher intensity. The amplitude of the estimated diffusion coefficient corresponds to the number of tracked particles.

Here we would like to point out that full capsids are detectable in larger detection volume than empty capsids because of their higher refractive index and the correlated larger signal to noise ratio allows us to probe particles further from the focus; indeed, despite the use of a similar concentration and that both populations have similar mobility, we have been able to track more than two times full capsids than empty capsids of T5. Therefore, for the estimation of nanoparticles concentration by SPT analysis, we should consider the influence of the particle size on the interferometric signal strength to normalize the volume of the detection.

### 3.2 Monitoring of virus-IgG recognition using Brownian motion

Here, we used the specificity of virus-antibodies reaction to detect significant modification using our set-up. We first dipped the objective lens in droplet containing purified polyclonal IgG molecules anti pb8 (major protein of the T5 capsid), and we then added T5 phage to the IgG solution. The T5 phage has about 800 binding sites, molecules of pb8 per capsid. We used an excess ratio of 2000 for each virus in the sample solution.

The measurements are performed in a binary logarithmic scale for the three samples of: the phages only; the phages added the antibodies solution and the antibodies only. **Fig 4** (a) and (b) processed images of the control T5-phages and the T5 phages in the presence of specific antibodies at different time intervals, respectively. A visual inspection of the T5 control does not reveal any significant differences as a function of time. However, a difference in terms of contrast, number of spots and size of the detected spots could be visualized after adding T5 phages to antibodies solution. This modification is confirmed using single-particle tracking as shown in **Fig 4** (c-d). A progressive decrease of the diffusion coefficient is observed for the sample containing the IgG molecules and T5 phages. A Diffusion coefficient of 3.11 *µm*^2^/*s* has been estimated 1 minute after adding T5 phages to IgG solution, this value gradually decreases to reach 1.17 *µm*^2^/*s* after 64 minutes. Based on the Stokes-Einstein equation, the hydrodynamic radius of individual particles increases by a factor of 3 in 64 minutes, (see **Fig 4** (c)). The modification of the diffusivity of the particles is paralleled the continuous rise of the interferometric signal as shown in **fig 4** (d). To note that we have not detected a significant modification for the T5 control in terms of diffusion coefficient, interferometric intensity in **Fig** 4 (c,d)

**Fig 4:**
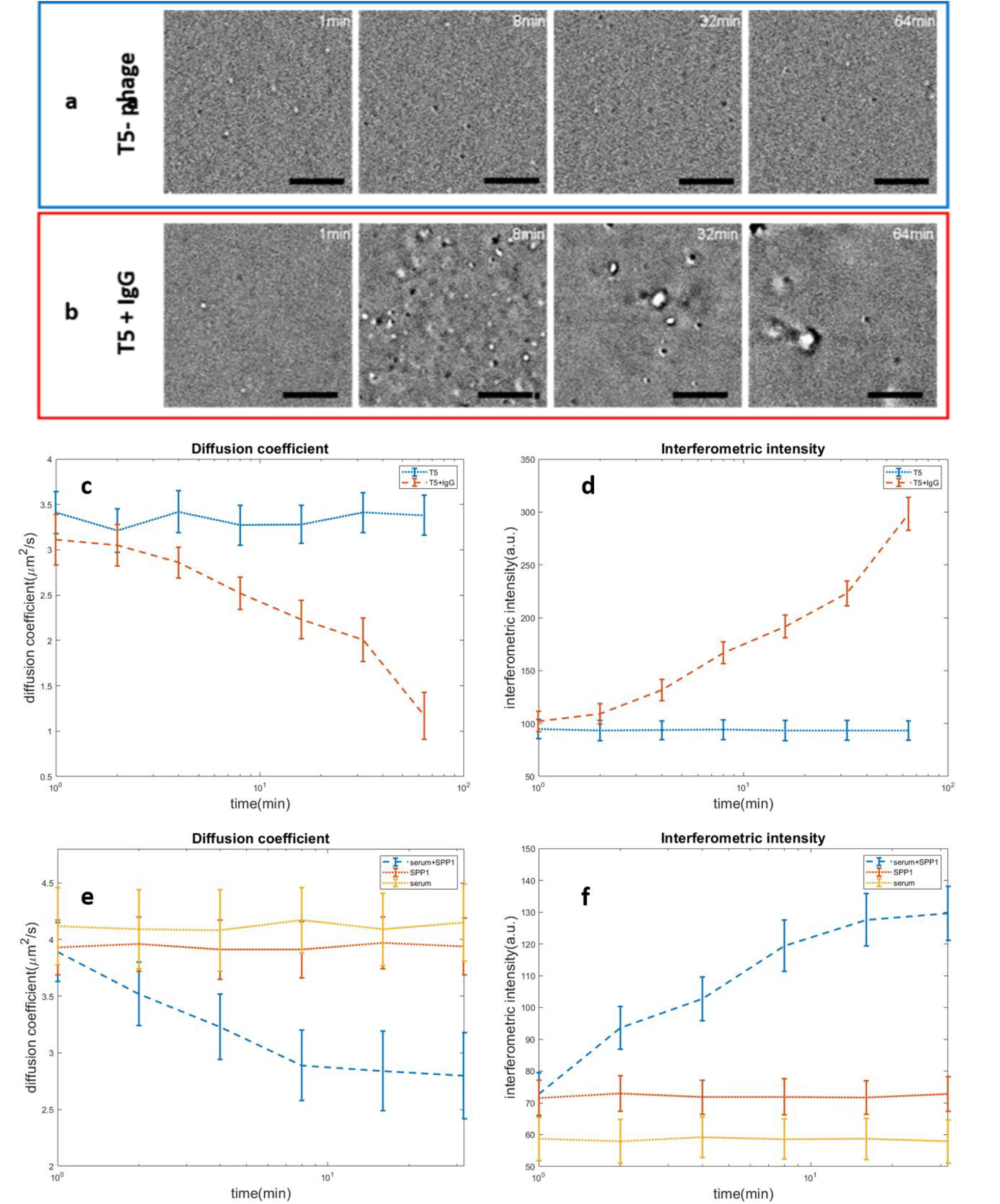
(a) Snapshots at different lag of time of the T5 bacteriophages alone and (b) the interaction in presence of IgG anti pb8 at different lag of time, respectively. (c), and(d) represent the diffusion coefficient andthe maximum of intensity of T5, T5+IgG respectively as function of time. (e) and (f) represent the diffusion coefficient and the maximum of intensity respectively of Spp1 +serum, the Spp1 and serum control as function of time.

The increase of the hydrodynamic radius and the interferometric intensity confirms the aggregation of the virus particles in the presence of antibodies.

We would like to mention that although the full capsid of T5 and the T5 phages have similar capsid structures, a difference in diffusion coefficient has been detected between both particles. The main difference affecting the dynamic behavior of these particles is the presence of a flexible tail used by the phage to infect its host. We would also note that we did not find significant modifications of intensity, diffusion mobility, and the number of particles for the control test of IgG molecules only and a sample of T5 added to purified antibodies targeting VSV virus (see supplementary). The movie of the interferometric images of T5 + anti pb8 at different time intervals in <u>Visualization 1</u> in the supplementary document.

### 3.3 Monitoring the serum-viruses reaction

We demonstrated the advantage of using the Brownian motion to follow the interaction between specific phage and adding purified antibodies. Here, we expand the application of this approach by following the interactions between SPP1 phage and non-purified serum containing multiple polyclonal antibodies molecules targeting different protein presented on the outer surface of the virus, neither on the tail or the capsid of the phage. Serum can be more challenging since it consists of different types of particles that do not necessarily interfere with the interaction between antibodies and the virus, these nanoparticles could affect single particle tracking analysis. The use of serum can be beneficial since it represents real physiological conditions for usual biological samples. For our measurements, we employed a final concentration of a final concentration of 2 × 10^9^*PFU/mL* of SPP1 phages in the serum solution.

A progressive decrease of the diffusion mobility of individual particles has been detected for the sample containing SPP1 phages and the anti-SPP1 serum. The diffusion coefficient reduce from 3.89 to 2.79 *µm*^2^/*s* in 32 minutes, **Figure 4** (e). This increase of the average size of the particles is confirmed by the rise of the interferometric signal, we observed a gradual rise of the signal from 72.8 to 129.6 (*a. u*.) in 32 minutes, **Figure 4** (f). We would like to point that as shown in the **Figure 4** (e, f), the diffusion coefficient and the interferometric signal, do not vary as function of time for the control sample of the SPP1 and the serum sample only. These results allows us to identify and follow the reaction between the phages and serum targeted virus. Despite that the serum contains various types of particles in addition to multiple antibodies molecules, the reaction between the serum and targeted phage has been detected in few minutes.

### 3.4 Quantitative analysis of the IgG-T5 reaction

Using single particle tracking microscopy and our MSD temporal evolution analysis, we introduce a quantitative approach to estimate the dissociation rate of specific antibodies. Here, we monitor the variation of the aggregation process depending on the initial concentration of antibodies in the solution sample.

As mentioned above, single antibodies cannot be detected by the optical set-up and the aggregation between viruses and antibodies is not homogeneous. Hence, we rely on the number of tracked viruses only (particles having a similar diffusivity as the T5 control sample). We therefore estimate the number of tracked viruses for the T5 control parameters in terms of the diffusion coefficient and counts of tracked particles, **Fig 5** (a). Then, we compare the diffusion coefficient histogram curve for different concentration of antibodies and for different time interval to the mean diffusion coefficient curve of the T5 control sample. When adding the antibodies to the virus solutions, the number of tracked particles varies for different time interval. Therefore, by normalizing the diffusion coefficient curves and comparing it to the control samples, we estimate a percentage of virus particles ratio. The number of virus particle is estimated using the total number of tracked particles. **Fig5 (b)** shows the diffusion coefficient histogram curve 1 min after adding the antibodies, for a concentration of 0.042 mg/mL, that corresponds to 2.8*10^−7^M. When comparing the diffusion coefficient histogram curve 64 minutes after adding antibodies, a clear shift towards smaller diffusion coefficient can be seen (**fig5 (c))**. Based on the estimated ratio resulting from the diffusivity comparison, we can estimate the number of viruses not yet interacting with antibodies using the number of total tracked particles for different dilutions of Ab as shown in **Fig5 (d)**. Hence, we estimate the number of bound viruses by subtracting the number of remaining viruses from the initial number of viruses.

**Fig 5:**
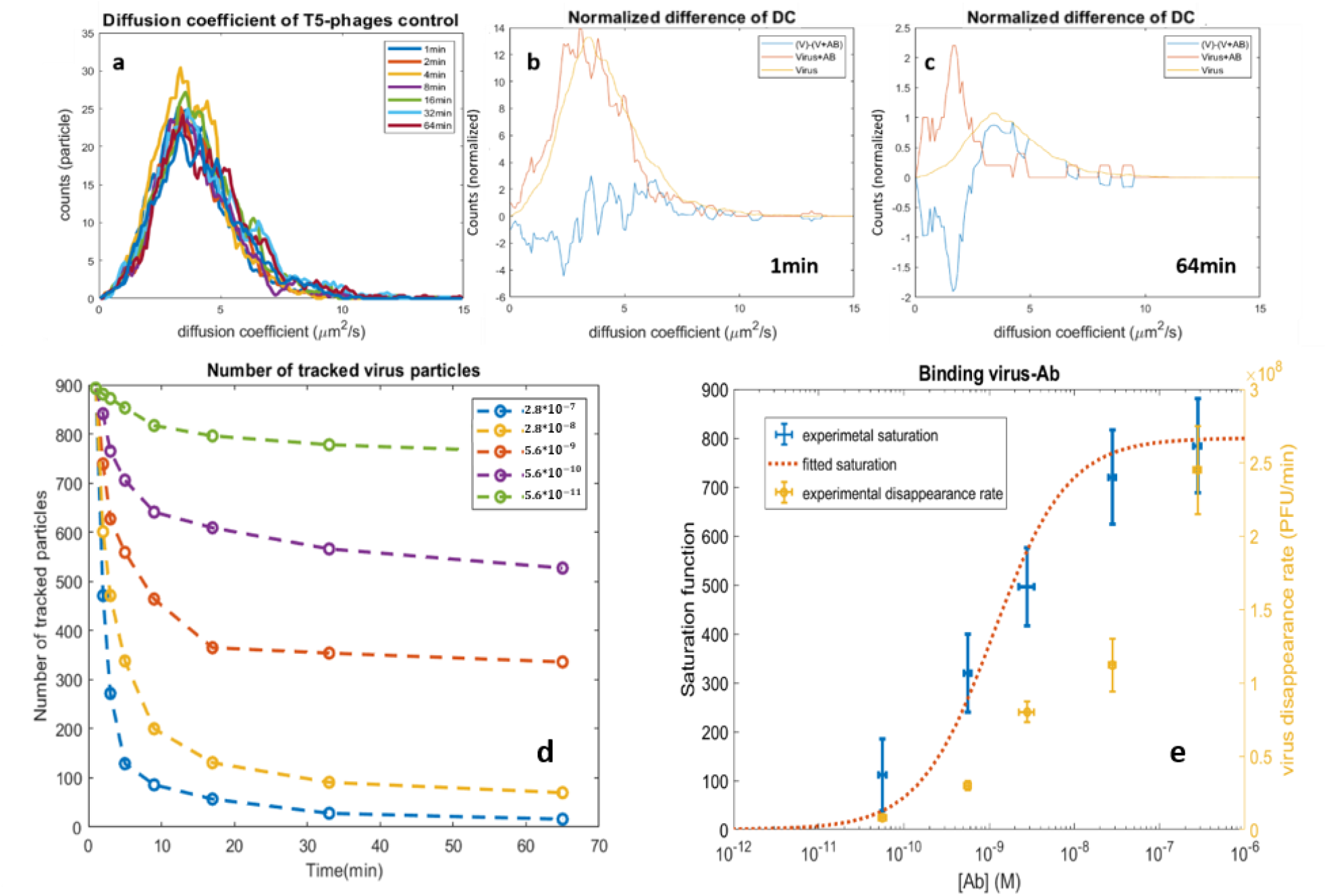
(a) Diffusion coefficient histogram curve of T5 phage control for different interval of time. (b) and (c) represent the normalized diffusion coefficient histogram curve for the control of T5 and T5 + Ab as well as the difference between the two curves for an interval of 1 and 64min respectively. (d) shows the number of virus particles tracked in the sample solution at different time interval when varying the concentration of antibodies, the legend of the plot indicates the concentration of antibodies in **M**. (e) represents the curve of the saturation function of the binding reaction on the left axis and the right axis represents the disappearance rate of the tracked virus particles as function of the concentration of An in the sample solution.

**Fig5** (d). shows two behaviors of the number of tracked virus particles, specifically for high concentration of antibodies. For small time interval, the number of detected viruses decrease rapidly in few minutes. Then, it reaches an asymptotic behavior indicating that the system reached its chemical equilibrium. Using the data points at 64min for different concentration, we estimate the number of bound antibodies in the solutions based on the estimation that all the tracked non virus particles have free binding sites, as for pb8 capsid protein, n=800 binding sites. As presented in the material and method section, the saturation function can be estimated based on the concentration ratio of viruses and antibodies in the solution sample. Fig5 (e) shows on the left axis the saturation function of the binding as function of the concentration of Ab. Thus, based on equation (6,b) by fitting the experimental data using the saturation function expression, we estimate ***K***_*d*_ = 1.12 **\*** 10^−9^*M*.

Furthermore, the reaction between phages and antibodies can be described on a higher scale, we are interested in measuring the speed of the reaction depending on the concentration of Ab in the solution. We attribute a half-life before reaching the system’s equilibrium state when varying the initial concentration of Ab. Despite the equilibrium state being reached at a different time interval, we attribute the number of virus particles at 64 minutes to the equilibrium. Then, we estimate the variation of the number of virus particles of the time interval corresponding to the half-life time ([*V*]_0_ − [*V*]_64_)/2. The right axis in **fig** 5(e) shows the disappearance rate of virus particles in the solution. In order to know the equation rate of the reaction, the proportional coefficient based on the stoichiometric coefficient is needed. Although we plotted the disappearance rate of viruses in the solution as a function of the concentration of antibodies, despite that this disappearance rate of viruses is proportional to the rate equation, we still need to find the stochiometric ratio of the reaction to be able to retrieve the rate equation of the reaction.

## 4. CONCLUSION

Through these examples, we have combined the interferometric signal contrast to the single particle tracking analysis to analyze the diffusion mobility and the component of the particles. This approach has been used to distinguish between two populations of capsids that have a similar size. Furthermore, viruses have been added to specific antibodies in order to monitor the antibody-virus reaction evolution in a wide field of view (103*103*µm*^2^), the recognition signature has been detected in few minutes. We also demonstrate the application of this approach to monitor the evolution of the reaction between phages and non-purified serum, which represent more realistic way to study physiological samples of viruses. However, further technical improvement can be made to reach better sensitivity and sensibility.

We have shown in this study that a simple, robust incoherent illuminated common path interferometer can be used to characterize nanoparticles in biological samples. This label-free detection method has been used to detect a specific reaction between viruses and specific antibodies in solution without complicated processing. Here we have used the specificity using the antibodies and non-purified serum by monitoring a modification in terms of particles mobility and the interferometric signal contrast. The simplicity of the detection method and the use of antibodies for specificity and processing might define new perspectives for virus and antibodies detection as a medical diagnosis tool and a control tool for the pharmaceutical drug delivery methods using viruses and vesicles. Such simple, robust and easy to use approach consists an efficient way to estimate the presence and concentration of antibodies for diagnosis applications as well as detecting the neutralization of viruses when adding specific antibodies.

Further improvements can be considered to increase the sensitivity of the interferometric set-up by attenuating the reference beam to match the weak scattered light beam coming from the particles [28]. One way to perform such attenuation of the reference beam is to block partially the incident light in the back focal plane of the objective lens [29]. Although that the increase of the interferometric contrast increase the sensitivity as well as the sensibility of the system, it requires the use of more complicated approach than presented in this work.

## Supporting information

the supplemental fig1 and fig3 show SPT and the statistical analysis for SiO2, 100nm. Fig 3 shows a negative control of T5+ anti VSV

---

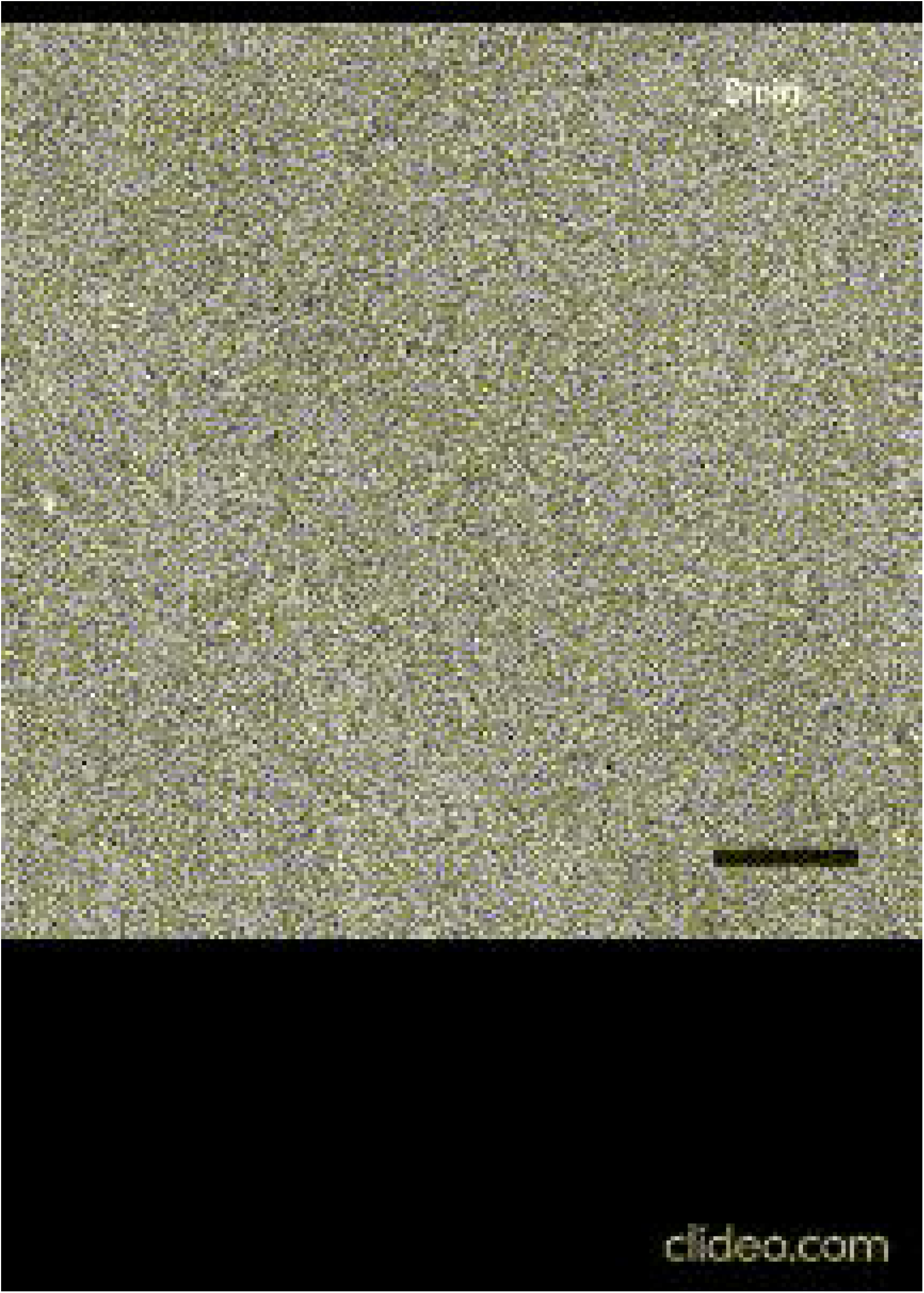

