## Supplementary material for "Label-free virus-antibody interaction monitoring in real time by common-path interferometry": the supplemental fig1 and fig3 show SPT and the statistical analysis for SiO2, 100nm. Fig 3 shows a negative control of T5+ anti VSV

***[1] Institut Langevin, ESPCI Paris, CNRS, PSL University. 1 rue Jussieu, 75005 Paris***
***[2] Institut de Biologie Intégrative de la Cellule, Université Paris-Saclay, CNRS, CEA, UMR9198, Orsay***
***[3] Institut de Biologie de l'ENS, CNRS UMR8197, Inserm U1024. 46 rue d'Ulm, 75005 Paris***

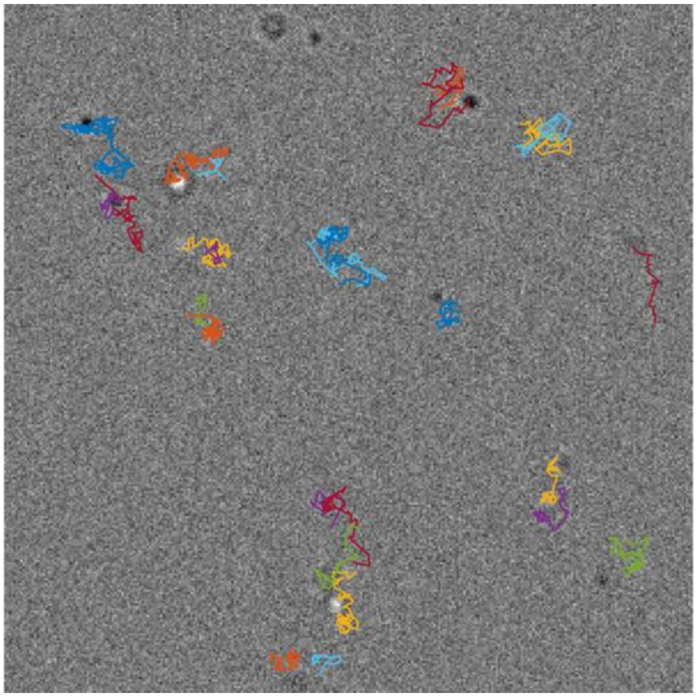

Fig1: Single particle trajectories resulting from the modified version of MTT software

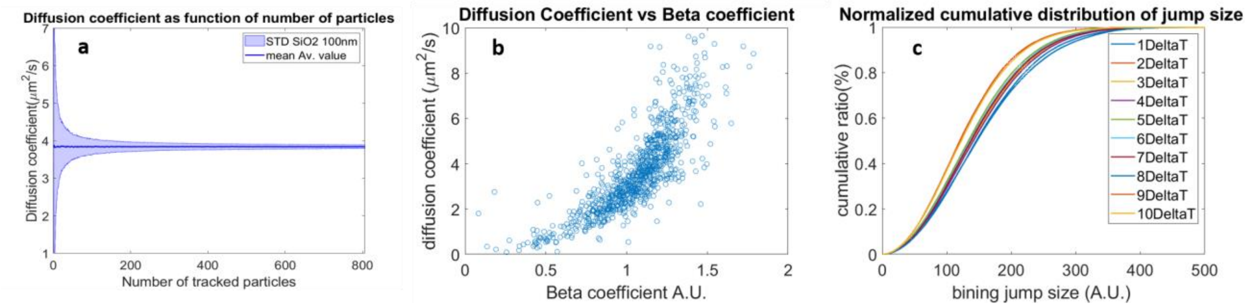


Fig2: Statistical representation of the single particle tracking analysis. (a) shows the variation of the mean estimated diffusion coefficient of SiO2 nanoparticle of 100nm in diameter. (b) represents the estimated diffusion coefficient and the beta coefficient for the single tracked particles. (c) shows the cumulative probability distribution of the jump size for different lag of time to check the random motion characteristic of the particles.

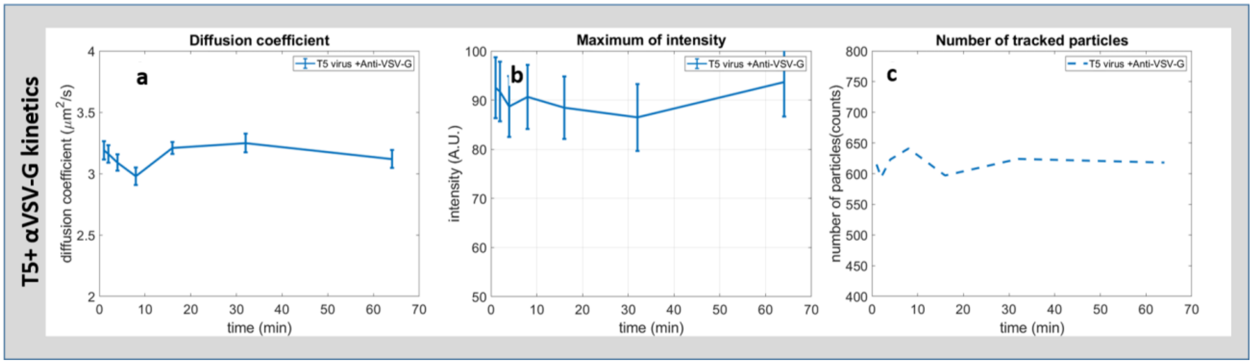


Fig3: evolution of T5 phage in presence of antibodies anti VSV. (a) plot representing the variation of the diffusion coefficient. Representation of the evolution of the interferometric intensity as function of time. The variation of the number of tracked particles as function of time.
